# Disentangling the spread dynamics of insect invasions using spatial networks

**DOI:** 10.1101/2022.12.15.520585

**Authors:** Sergio A. Estay, Carmen P. Silva, Daniela N. López, Fabio A. Labra

## Abstract

Describing and understanding spatiotemporal spread patterns in invasive species remains a longstanding interdisciplinary research goal. Here we show how a network-based top-down approach allows the efficient description of the ongoing invasion by *Drosophila suzukii* in Chile. To do so, we apply theoretical graph methods to calculate the minimum cost arborescence graph (MCA) to reconstruct and understand the invasion dynamics of *D. suzukii* since the first detection in 2017. This method estimates a directed rooted weighted graph by minimizing the total length of the resulting graph. To describe the temporal pattern of spread, we estimate three metrics of spread: the median dispersal rate, the median coefficient of diffusion, and the median dispersal acceleration. The estimated MCA shows that over four years, *D. suzukii* colonized a ~1,000 km long strip in the central valley of Chile, with an initial phase with long paths and connections and no clear direction pattern, folowed by a clearer north-east propagation pattern. The median dispersal rate for the entire period was 8.8 [7.4 – 10.6,95% CI], while the median diffusion coefficient was 19.6 meters^2^/day [13.6 – 27.9, 95% CI]. The observed spread dynamics and the log-normal distribution of accelerations are consistent with longdistance dispersal events. The complexities of real landscapes cannot be summarized in any model, but this study shows how an alternative top-down approach based on graph theory can facilitate the ecological analysis of the spread of an invasive species in a new territory.

## Introduction

Describing the spatial spread in invasive species and understanding the underlying processes across different organisms have been long-standing research goals across many disciplines ranging from biology to mathematics and epidemiology (Fisher, 1937; Skellam, 1951; Andow, et al. 1990; Okubo and Levin, 2002; Hastings et al., 2005). Among the stages of an invasion process, spread corresponds to invading organisms moving across the landscape and colonizing new habitats. Spread is a critical phase in the invasion process froma biological point of view (e.g., Andow et al., 1990) and control and management perspective (Epanchin-Niell et al., 2010; Epanchin-Niell and Wilen, 2012; Robertson et al., 2020). Based on the early works of Fisher (1937) and Skellam (1951), most work on spread modeling has followed a mechanistic bottom-up approach (Hordijkand Broennimann, 2012), using integrodifference or reaction-diffusion models combined with a population growth model (Van den Bosch et al., 1990; Kot et al., 1996). This approach has the advantage of providing an explicit representation of the different processes involved in the invasion process (Shigesada and Kawasaki, 1997). It allows the estimation of spread descriptors by fitting models with mathematically explicit functional forms and parameters. However, spread is a complex process, and the assumptions of these models could be too simplistic, making it difficult to obtain robust reconstructions of the invasion dynamics and the rate of spread through space and time (Hastings et al., 2005; Hordijk and Broennimann, 2012). In addition, most mechanistic bottom-up invasion spread models are usually defined using a deterministic framework (Kot et al., 1996; Lewis, 2000; Lewis and Pacala, 2000), ignoring the importance of extrinsic or intrinsic stochastic factors (Hastings et al., 2005). Extrinsic stochasticity sources may include the host’s actual spatial distribution, the climate effect on insect phenology, or the natural barrier’s presence (Carrasco et al., 2010). Intrinsic sources of stochasticity may include the exact form of the dispersal kernel or stochastic population dynamics (Lewis, 2000). On the other hand, empirical approaches to the description of invasion processes and invasion spread emphasize the role of spatial heterogeneity, temporal variability, and ecological interactions (Hastings et al., 2005). These more versatile top-down approaches depend on a few assumptions, facilitatating their implementation (Hordijkand Broennimann, 2012). Thus, top-down approaches may provide spread descriptors without assuming specific mechanisms or explicit mathematical models (Hordijk and Broennimann, 2012), which may be argued to be their major disadvantage.

In the top-down approach, models based on network theory have successfully been applied to describe the spread phenomenon. Disease propagation is a special case of a spreading phenomenon studied using networks (e.g., Newman, 2002; Sattenspiel and Lloyd, 2009; Björnstad, 2018). In this case, nodes in the network may be either people or sites (e.g., cities), and links represent contacts or connections (routes). In these spatial networks of sites or cities, the routes are defined a priori (e.g., highways), and spread occurs through these links. However, a slightly different approach exists to analyzing invasive species spread dynamics. This approach describes the geometrical properties of species invasion patterns by using graphs such as the Minimal Spanning Tree (MST) (Labra et al., 2005). But also describes the invasion process by successive invaded sites or sampling areas where an invasive species has been recorded. These invaded sites are represented in the two-dimensional geographical space as a set of nodes, with the Euclidean distance describing the geographical separation among them (Fig. 1a). These points in space may be joined in many different ways by lines (or edges), forming a graph. The resulting graph may be defined as connected if there is an edge between any pair of nodes, as shown in Fig. 1b, which includes several circuits or loops (Labra et al., 2005). It has been shown that for complex spatial patterns, the dominant pattern of connectedness may be described by a minimal spanning tree (MST), which is a graph that connects all the nodes with no circuits or loops. It only considers those connections that minimize the total length across the graph (Fig. 1c) (Prim, 1957). MST has a long history of application in biology and ecology to describe various spatial patterns (Dussert et al., 1986, 1987; Cantwell and Forman, 1993; Lockwood et al., 1993; Jones et al., 1996; Keitt et al., 1997; Wallet and Dussert, 1997; Bunn et al., 2000, Urban et al., 2001; Labra et al., 2005). However, to fully describe the invasion process, the MST should be rooted at the first recorded invasion site and directed to reflect the temporal sequence of site invasion. Optimum branching (OB), minimum cost arborescence (MCA, Fig. 1d), or directed minimum spanning tree (DMST) are all names that have been used to describe same problem.Given a directed rooted weighted graph, what is the minimum cost (sum of weights) graphwith a unique path from the root to any other node? (Chu and Liu, 1965; Edmonds, 1967; Tarjan, 1977). In the context of the spread of an invasive species, if the original point of introduction and successive dated points where the species was detected are known, and if we assume that spread follows the shortest path, then we can use the minimum cost arborescence to reconstruct the spread of the species in the landscape. The resulting MCA is a directed weighted graph with weights equal to the Euclidean distance between nodes (sites), the in-degree for all nodes is always one, as it is a branching process where new sites are assumed to originate from a single previously invaded site. However, the out-degree can show some variability depending on the topology of the resulting network, as any given site may be linked to one or more newly invaded sites in the following time step.

**Figure 1:**
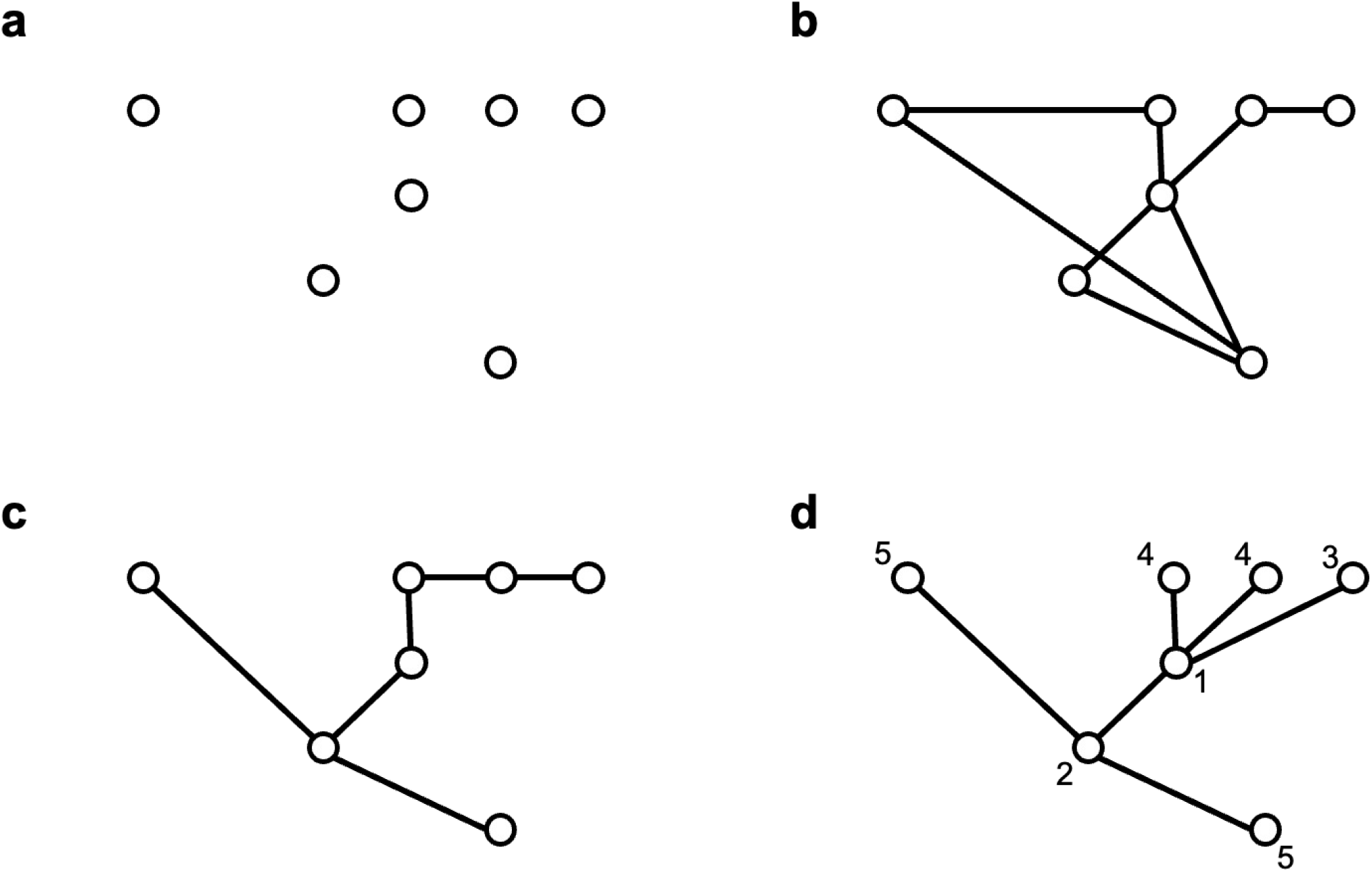
Examples of point and graph depictions of an invasive process. The figure shows (a) a set of successive invasion points in space, reflecting a number of iterations in an invasive spread process. (b) a connected line graph which connects the invasive points, showing sone loops. (c) a minimal spanning tree (MST) which links all the invaded points in a way to minimize the total Euclidean distance span of the tree. (d) an optimal branching tree or minimum cost arborescence graph which includes the information on the sequence of invasion of the different points in space, which is shown by the labels next to each data point.

Hordijk and Broennimann (2012) provide an excellent description of this approach; however, scarcely explored in the literature despite its many advantages. Among the advantages of this approach are that it facilitates the estimation of dispersal rate through the calculation of spread velocity and acceleration and allows the detection of long-distance jumps. In addition, metrics show changes in response to spatiotemporal heterogeneity, and hence may be valuable to identify different environmental drivers that may be forcing or influencing the invasive spread. Here we show how this approach may be used to describe an ongoing biological invasion.

Exotic insect pests colonizing new ecosystems have caused multiple detrimental effects around the world, having a direct impact on agricultural and forestry production (e.g., Bradshaw et al., 2016), ecosystems services (e.g., Clark et al., 2010), human health (Lounibos, 2002), and cultural values (Manachini, 2015). Among the most important insect pests currently attacking fruit production worldwide is *Drosophila suzukii.* This species originated in South East Asia, but in 2008, it was detected simultaneously in the USA, Italy, and Spain (Rota-Stabelli, 2013). Since then, it has been introduced in several regions like Hawaii, North America, Europe, Central America, South America, and Africa (Rota-Stabelli, 2013; CABI, 2022). Adults of this species are 2-3 mm long with red eyes, pale brown or yellowish-brown thorax, and black transverse stripes on the abdomen (CABI, 2022). *Drosophila suzukii* shows a winter morph with increased cold tolerance and larger wings (Shearer et al., 2016). In 2017, *D. suzukii* was detected for the first time in Chile (Rojas et al., 2019; Devotto 2020), and in the following years, the species spread through the country. Berries are among the most important fruit host of *D. suzukii.* In 2020 Chile had ~22,000 ha planted with these species (Garcia et al., 2022). Buzetti (2020) reported average direct damage per year between 1.2 and 2.7 tonnes/ha with losses of 5,000 to 17,550 US$/ha in cherries. In blueberries, damage ranges between 1 and 1.5 tonnes/ ha, equivalent to 4,000 US$/ha (Buzzetti, 2020). After the first detection, the Agricultural and Livestock Service (SAG in Spanish) implemented an intensive monitoring program that continues until today, where daily georeferenced captures are registered and reported (SAG, 2021). Currently, this is an ongoing invasion; hence the estimation of the spread rate is a fundamental metric with significant economic and management implications. In this study, we apply MCA methods to reconstruct and understand the invasion dynamics of *D. suzukii* in Chile since the first detection in 2017. We used this graph-based approach to address the following research questions: 1) what is the magnitude of spread? 2) how does the rate of spread vary when different periods and geographic regions are compared? 3) What is the relationship between the spatial and temporal variation and the spread rate with the species biology? Our results will contribute to understanding this species’s invasion biology and could provide valuable tools for developing appropriate control measures.

## Methods

### Data

After the first detection of *D. suzukii* in Southern Chile in June 2017, the SAG implemented an intensive monitoring program based on traps and visual inspections (SAG, 2017). We obtain dated georeferenced data points (sites) from SAG’s open repository (shorturl.at/AKOR2). We used data collected between June 9th, 2017 to June 29th, 2021. We eliminated the first point (June 5th, 2017) because it is an isolated point and probably corresponds to a secondary dispersion. After June 29^th^, 2021, the expansion *D. suzukii* continues, but through very long-distance dispersal in the Northern, the arid region of Chile. These jumps are probably linked to long distance human transport, so we decide to exclude this data from our analysis. In total, we used 3,907 data points (Fig. 2). To produce our results in meters, we reprojected all coordinates to WGS 84, UTM 19S CRS.

**Figure 2:**
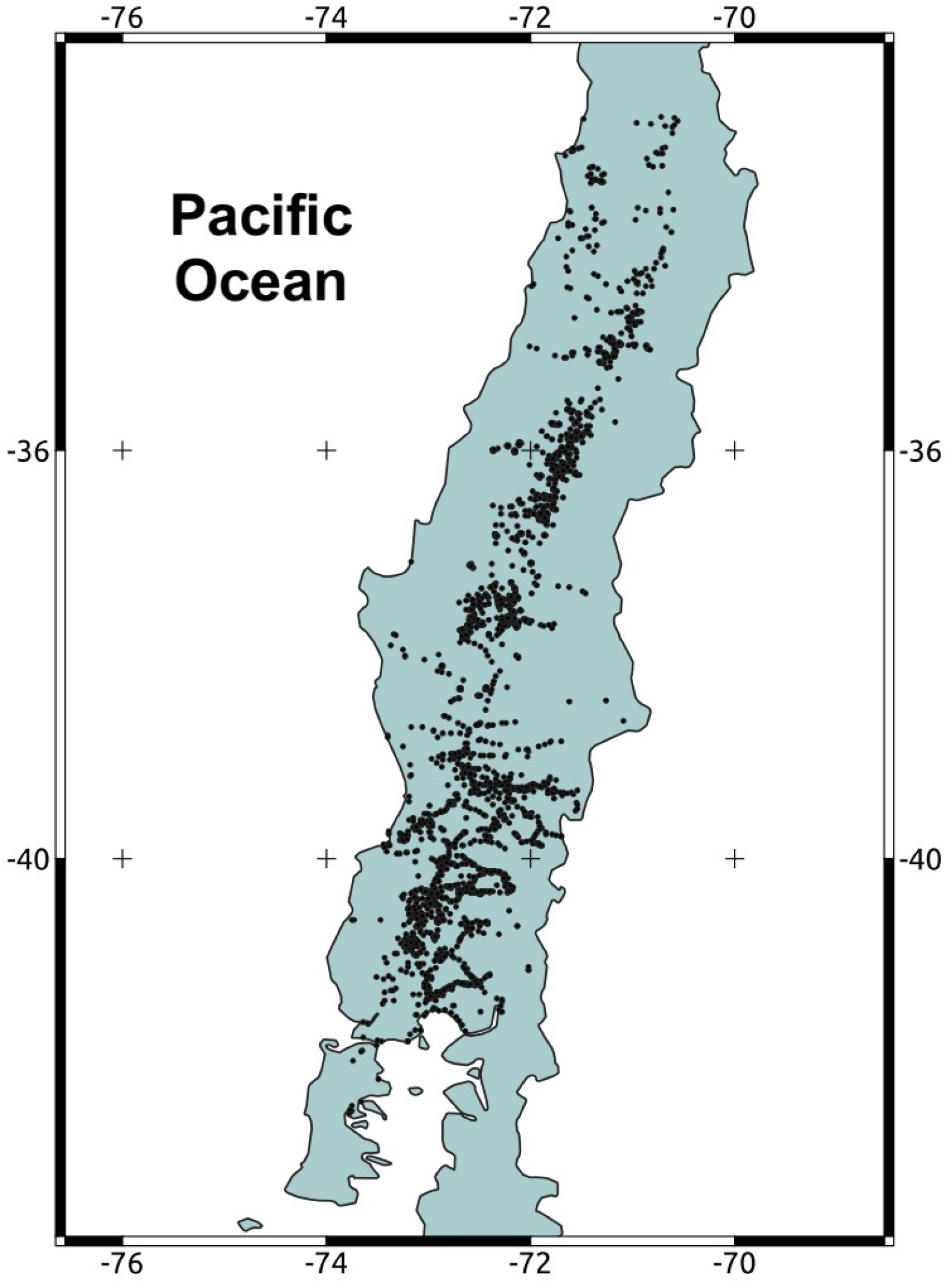
Map of the Chilean territory invaded by *Drosophila suzukii.* Dark points correspond to detections of the pest between the years 2017-2021.

### Analysis

Using the previously described data, we reconstruct the invasion dynamics of *D. suzukii* in Chile using MCA. Following Hordijk and Broennimann (2012), we rooted our MCA at the first detection point (first node), and the link weights between nodes correspond to the Euclidean distance. In our MCA, nodes correspond to sites positive to *D. suzukii,* and links correspond to the shortest path between nodes. We estimate one MCA for the complete period.

After estimating our MCA, we calculated several metrics to understand the spread dynamics. First, we estimated the median dispersal rate (Link weight/days) and the 95% confidence interval by bootstrapping the final results obtained from the MCA. We take 1000 pseudo-samples over 1000 iterations. To evaluate the temporal variability in dispersal rate, we calculate the median value over a moving window of 100 days (~three months). Second, to make our results comparable with other studies, we calculate the variability in the median coefficient of diffusion through time using 100-days moving windows. To do this, we use the estimated median values of dispersal rates and the formula described by Shigesada and Kawasaki (1997). Finally, we use the MCA methodology to estimate the variability in acceleration, calculated as the difference in dispersal rate through the study period. All calculations were performed in the R environment (R core team, 2022) using the package ecospat (Broennimann et al., 2022).

## Results

Between 2017 and 2021, *D. suzukii* spread rapidly throughout the country. In four years, the species has colonized a ~1,000 km long strip in the central valley of Chile, ranging from 32° to 42° Lat S (Fig. 2). The estimated MCA (Fig. 3) shows an initial phase with long paths and connections without a clear direction pattern (Fig. 3). After this phase, a clearer north-east propagation pattern emerges (Fig. 3). The median dispersal rate for the entire period was 8.8 meters/day with a 95% CI of [7.4 – 10.6] (Fig. 4a). The coefficient of diffusion showed a median value of 19.6 meters^2^/day with a 95% CI [13.6 – 27.9] (Fig. 4b).

**Figure 3:**
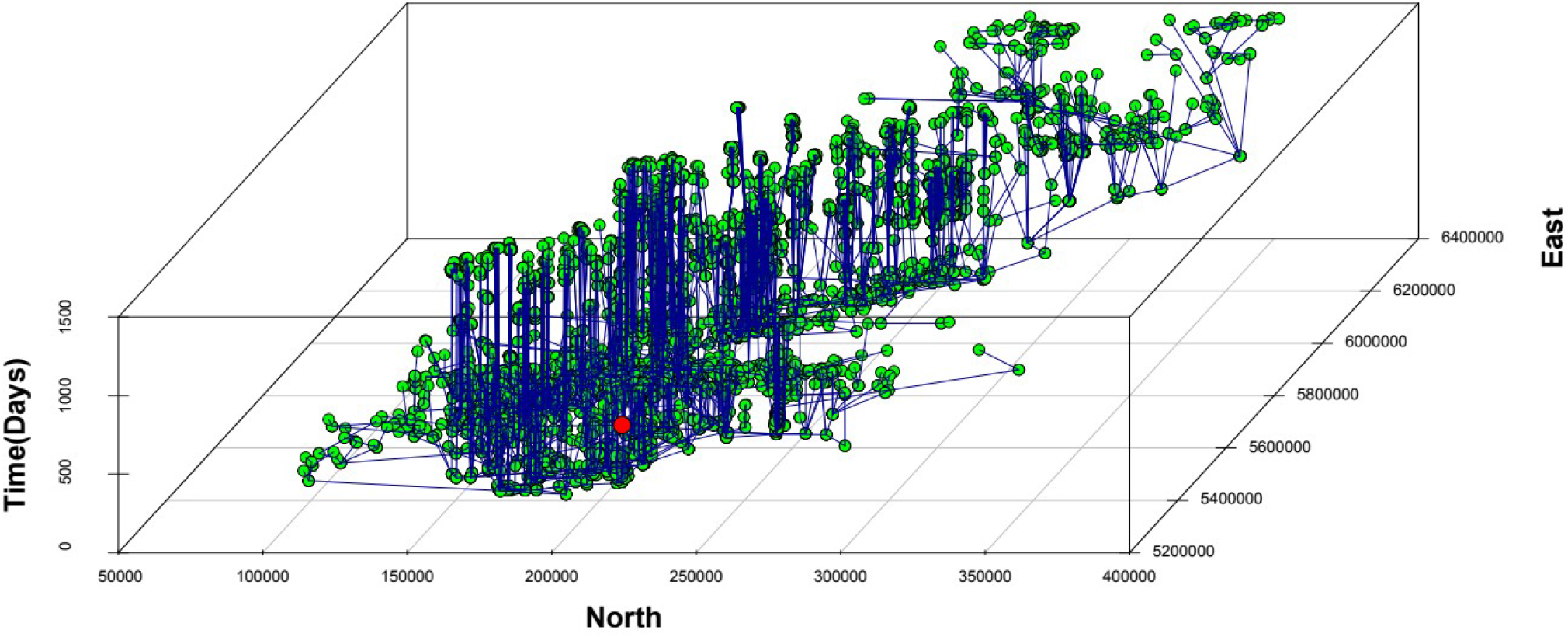
MCA reconstructing the invasion of *D. suzukii* in Chile. Green circles (nodes) correspond to sites, and black lines (links) represent the hypothetical spread paths between sites under the assumptions of the MCA in the three-dimensional space (north-south-time).The red circle marks our MCA’s root (initial point of introduction).

**Figure 4:**
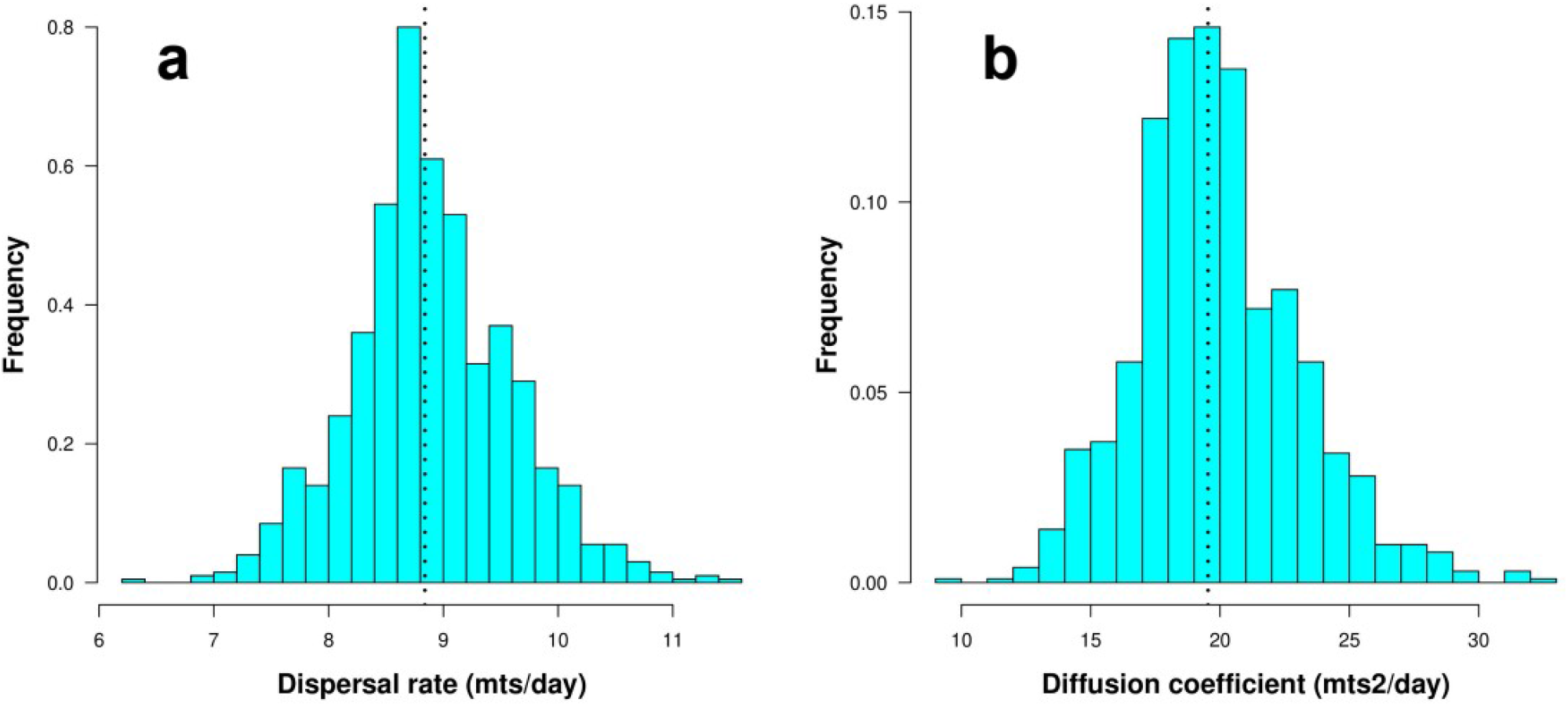
Histograms showing the distribution of values obtained through bootstrap. The dotted line corresponds to the median. a) Dispersal rate, b) Coefficient of diffusion.

A temporal look at the pattern of propagation shows two clear peaks of spread during the autumn and winter of 2017 and 2018, where 75% of the final area in Chile was reached. During the autumn-winter of the first two years, dispersal rates reached a maximum of ~110 meters/day, and the diffusion coefficient reached a maximum of 3,000 meters2/day (Fig. 5a-b). Both values are several times higher than the median values for the entire period. In a similar way, spread shows several acceleration-deceleration phases, also mainly in the autumn-winter season (Fig. 5c). After the first two years, the dispersal rate stabilizes around the median values. The dispersal rates follow a log-normal distribution, typical of processes that involve multiple scales, as can be seen when we compare the empirical probabilities with the theoretical probabilities from the best-fitting log-normal (Fig. 5d, see discussion).

**Figure 5:**
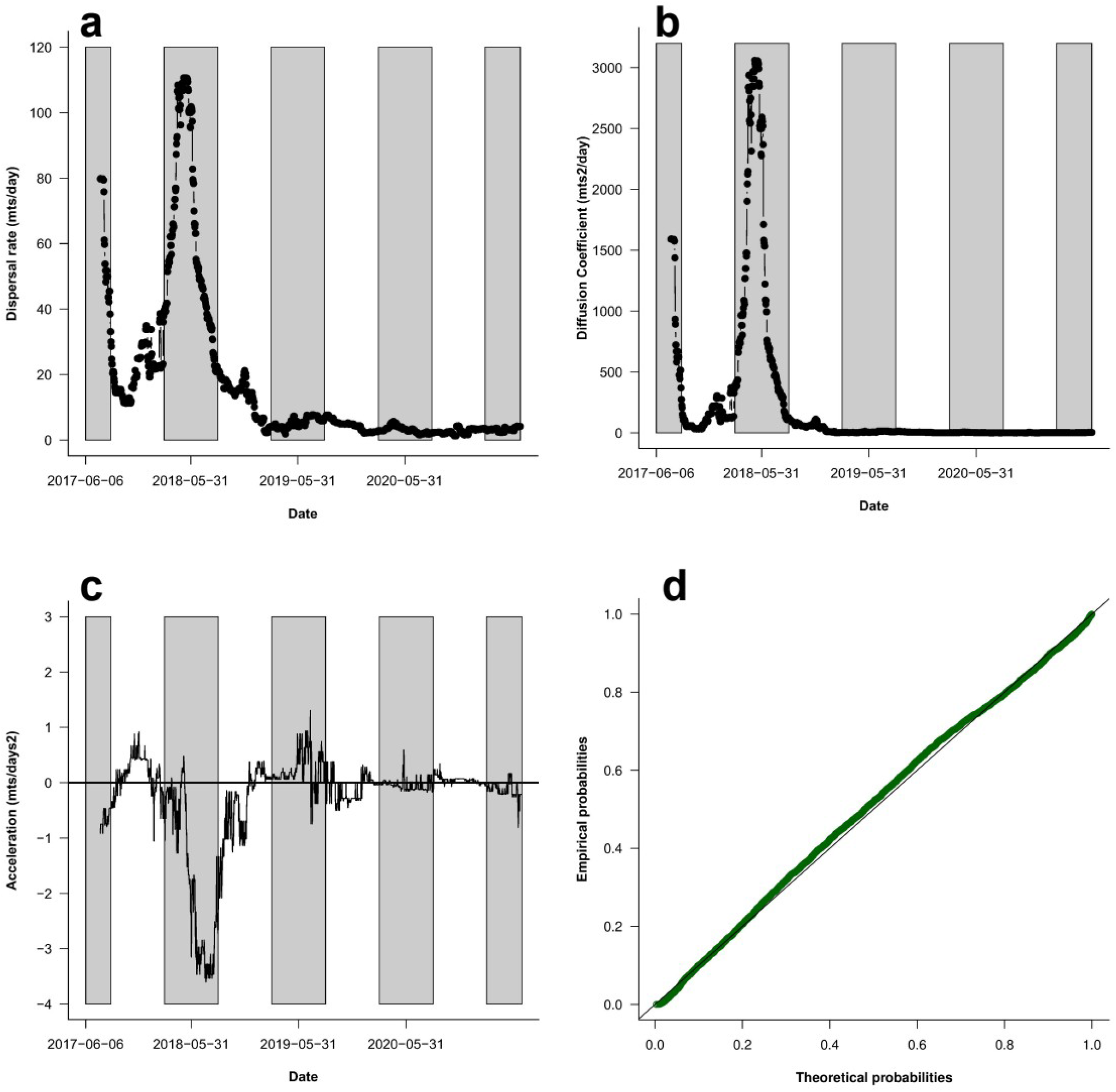
Temporal variation of the dispersal rate (a), coefficient of diffusion (b), and acceleration (c) through the study period using a 100-days moving window (see methods).Grey boxes correspond to the autumn-winter season of each year. d) p-p plot of the theoretical vs empirical values (green dots) of the log-normal distribution fitted to the data. The solid line corresponds to the 1:1 ratio (perfect fit).

## Discussion

*Drosophila suzukii* has spread rapidly along Chile, occupying most of the region with berries plantations in less than four years. This high spread capacity is a well-known trait of this species. Hauser (2011) and Calabria et al. (2012) have described the fast dispersal of *D. suzukii* in the USA and Europe, respectively. In particular, Calabria et al. (2012) pointed out a dispersal rate of 1,400 km/year, probably combining active and passive dispersal. In Chile, the species did not reach such an extremely high dispersal rate, but 1,000 km in 3 years is still a significantly high spread capacity. As it was suggested by Calabria et al. (2012), this high dispersal rate is the consequence of a combination of active (diffusion) and passive (human transport) spread, which is called stratified dispersal (Shigesada and Kawasaki, 1997). In our case, observed dispersal rates follow almost perfectly a log-normal distribution, a well-known heavy-tailed distribution. This result suggests that the occurrence of extreme, long-distance dispersal events is “more common” than that expected under diffusion-only or Gaussian spread dynamics, in line with the observation of Calabria et al. (2012).

The general estimation of 8.8 meters/day is close to the results obtained experimentally by Vacas et al. (2019). These authors, performing mark-release-recapture experiments, show that most individuals of the species were recaptured at a distance below 10 meters after 24 hrs. However, several studies have reported that *D. suzukii* is capable of long flights. Tait et al. (2018), also performing mark-recapture experiments, reported high variability in the dispersal rate of this species. Depending on elevation, these authors reported several individuals with almost no movements after several weeks of sampling, whereas other individuals show dispersal rates as high as 9,000 meters/month (~300 meters/day). In the same vein, Wong et al. (2018), described individuals moving over one km in a single flying event, which also supports the occurrence of long-distance dispersal events.

However, this median estimation can be misleading, as there is high variability in the dispersal rate of the insect depending on the seasons. Our results show two main peaks of dispersal in the first years after introduction. The dispersal rate during these peaks reaches 110 meters/day, more than ten times higher than the median value. These two peaks occurred during the first autumn-winter seasons. In this season, fruit availability is lower, which can promote longer flight events (Little et al., 2020). Also, in Chile adults show a characteristic winter morph, with increased cold tolerance and larger wings which may be interpreted as an intrinsic factor associated with higher dispersal rates (Shearer et al., 2016). However, Tran et al. (2022) indicate that the winter morph does not show a higher-flying performance than the summer morph, despite its apparent morphological advantages. In this regard, our data is clear in showing a faster and accelerated spread in the autumn-winter seasons, but the determination and testing of the exact ecological mechanism behind this rapid spread will require new studies. After the initial fast propagation during the first two years, dispersal rates remained close to the median values, without new peaks, probably due to the species reaching most of the suitable habitats. In our MCA, this can be inferred by the lower median dispersal rates and few new nodes (colonized sites) after the second autumn-winter season (Fig. 3).

On the other hand, extrinsic factors play an important role in modulating dispersal rates. For example, meteorological factors like temperature (Lantschner et al., 2014; Leach et al., 2019) or wind speed (Leitch et al., 2021) can be key for explaining the observed spread dynamics during an invasion. The high dispersal rates described by Tait et al. (2018) were observed from high to low elevations, which would suggest a major influence of winds in these events. This behavior has also been detected in other *Drosophila* species, where even longer wind-assisted jumps of ~12 km are possible (Leitch et al., 2021). In a more general analysis, Tait et al. (2020) showed that individual dispersion is a function of meteorological factors like temperature and humidity but also depends on the local diversity of alternative hosts. In this context, our results reinforce the complexities of the spread process and highlight the multiple difficulties of extracting detailed, ecologically meaningful information from real data. Intrinsic factors like population dynamics, phenology, or the emergence of seasonal morphs interact or depend on extrinsic forces like temperature, wind, or host availability, creating particular spread dynamics that can hardly be captured by one model, even in laboratory experiments (Melbourne and Hastings, 2009).

Our use of MCA allowed us to estimate several metrics of spread, successfully identifying both spatial and temporal variations. In this regard, the use of MCA, a rooted, directed weighted tree, shows several improvements over previously used MST, a minimal weighted tree method. In this regard, we show that MCA provides an efficient analytical process to describe observed invasions, which successfully identifies spatial and temporal heterogeneity in the observed rates of spread. By rooting the graph and requiring it to be directed, this approach integrates historical constraints. In addition, the use of minimal weights (similar to the MST) provides a set of parsimonious assumptions to describe the successive dispersal events from one time window to the next. While our data reflect a high-frequency standardized sampling effort, further research is needed to determine whether this method would still be successful with data captured at coarser temporal and spatial sampling grains. The complexities of real landscapes cannot be summarized in any model, but this study shows how an alternative top-down approach based on graph theory can facilitate the ecological analysis of the spread of an invasive species in a new territory.

## Acknowledgments

The authors were supported by ANID PIA/BASAL FB0002 and Fondecyt 1211114 to SAE, CPS and DNL and Fondecyt 1221153 to FAL.

